# Biasing the perception of spoken words with tACS

**DOI:** 10.1101/806059

**Authors:** Anne Kösem, Hans Rutger Bosker, Ole Jensen, Peter Hagoort, Lars Riecke

**Affiliations:** Lyon Neuroscience Research Center, Université Lyon 1, Lyon, France; Max Planck Institute for Psycholinguistics, Nijmegen, The Netherlands; Radboud University, Donders Institute for Brain, Cognition, and Behaviour, Nijmegen, The Netherlands; University of Birmingham, Centre for Human Brain Health, Birmingham, United Kingdom; Department of Cognitive Neuroscience, Faculty of Psychology and Neuroscience, Maastricht University, Oxfordlaan 55, 6229 EV Maastricht, The Netherlands

## Abstract

Recent neuroimaging evidence suggests that the frequency of entrained oscillations in auditory cortices influences the perceived duration of speech segments, impacting word perception (Kösem et al. 2018). We further tested the causal influence of neural entrainment frequency during speech processing, by manipulating entrainment with continuous transcranial alternating current stimulation (tACS) at distinct oscillatory frequencies (3 Hz and 5.5 Hz) above the auditory cortices. Dutch participants listened to speech and were asked to report their percept of a target Dutch word, which contained a vowel with an ambiguous duration. Target words were presented either in isolation (first experiment) or at the end of spoken sentences (second experiment). We predicted that the frequency of the tACS current would influence neural entrainment and therewith how speech is perceptually sampled, leading to a perceptual over- or underestimation of the vowel duration. Experiment 1 revealed no significant result. In contrast, results from experiment 2 showed a significant effect of tACS frequency on target word perception. Faster tACS lead to more long-vowel word percepts, in line with previous findings suggesting that neural oscillations are instrumental in the temporal processing of speech. The different results from the two experiments suggest that the impact of tACS is dependent on the sensory context. tACS may have a stronger effect on spoken word perception when the words are presented in a continuous stream of speech as compared to when they are isolated, potentially because prior (stimulus-induced) entrainment of brain oscillations might be a prerequisite for tACS to be effective.

## Introduction

Non-invasive transcranial alternating current stimulation (tACS) is an increasingly popular technique in auditory and language research (Riecke and Zoefel 2018; Zoefel and Davis 2017; Heimrath et al. 2016), with accumulating evidence showing that tACS efficiently affects sound processing and speech comprehension. Low-frequency tACS in the theta range (4 Hz) and alpha range (10 Hz) influences sound detection (Riecke et al. 2015; Riecke, Sack, and Schroeder 2015; Neuling et al. 2012) and high-frequency (40 Hz) tACS affects phoneme categorization (Rufener, Zaehle, et al. 2016; Rufener, Oechslin, et al. 2016). During continuous speech listening, tACS modifies auditory speech-evoked activity in the auditory cortex (Zoefel, Archer-Boyd, and Davis 2018) and speech comprehension (Riecke et al. 2018; Wilsch et al. 2018).

The effects of tACS on auditory perception are thought to be mediated by oscillatory neural mechanisms that would be critical for auditory and linguistic processing (Giraud and Poeppel 2012; Peelle and Davis 2012; Zoefel, ten Oever, and Sack 2018). Previous evidence shows that neural activity in the auditory cortices tracks the rhythmic structure of the speech signal. This neural tracking is linked to speech processing: neural tracking is stronger when sentences are intelligible (Peelle, Gross, and Davis 2013; Ding and Simon 2013) and indicates how the speech signal is parsed in the brain (Kösem et al. 2018; Ding et al. 2016; Ten Oever and Sack 2015). tACS is thought to influence neural tracking by modulating oscillatory activity of neural networks (Fröhlich and McCormick 2010; Witkowski et al. 2016; Thut, Schyns, and Gross 2011) (but see (Asamoah, Khatoun, and Mc Laughlin 2019)), and hence may provide a technique to test for a causal influence of neural tracking on the comprehension of spoken language.

So far, most tACS studies on speech have focused on effects of tACS phase, that is, how the temporal alignment of the tACS current and speech envelope affect speech comprehension. Here, we further investigated whether the *frequency* of tACS influences speech perception. Neural activity in the theta range (3-8 Hz) is known to flexibly follow the syllabic rate of ongoing speech (Ahissar et al. 2001; Kösem et al. 2018). The flexible tracking of speech could reflect neural entrainment mechanisms, i.e. the endogenous adjustment of neural rhythms to sensory dynamics (Obleser and Kayser 2019). Neural entrainment are thought to facilitate speech processing via temporal referencing and temporal prediction (Kösem and van Wassenhove 2017; Kösem, Gramfort, and van Wassenhove 2014). The frequency of entrained theta oscillations would then define the expected syllabic rate from a brain referential standpoint, and this would influence how syllabic units and their constitutive phonological segments are processed in time (Fig. 1B) (Bosker 2017; Bosker and Kösem 2017; Kösem and van Wassenhove 2017; Kösem et al. 2018; Bosker and Ghitza 2018). Recently, findings from a magnetoencephalography (MEG) experiment by Kösem et al. (2018) provide support for this proposal. They show that sentences produced at a fast speech rate induce entrainment at a higher frequency (compared to slower sentences) and that this faster entrainment sustains for a few cycles after the driving stimulus has ceased. Moreover, this sustained entrainment influences behavioral categorization of subsequent ambiguous target words. This suggests that the neural tracking of the temporal dynamics of speech is a predictive mechanism that is involved in the processing of subsequent speech input and directly influences speech perception. In line with Kösem et al. (2018), we predicted in the present study that modulating the frequency of entrained theta oscillations with tACS modifies the perceived duration of speech segments and affects the perception of words.

**Figure 1:**
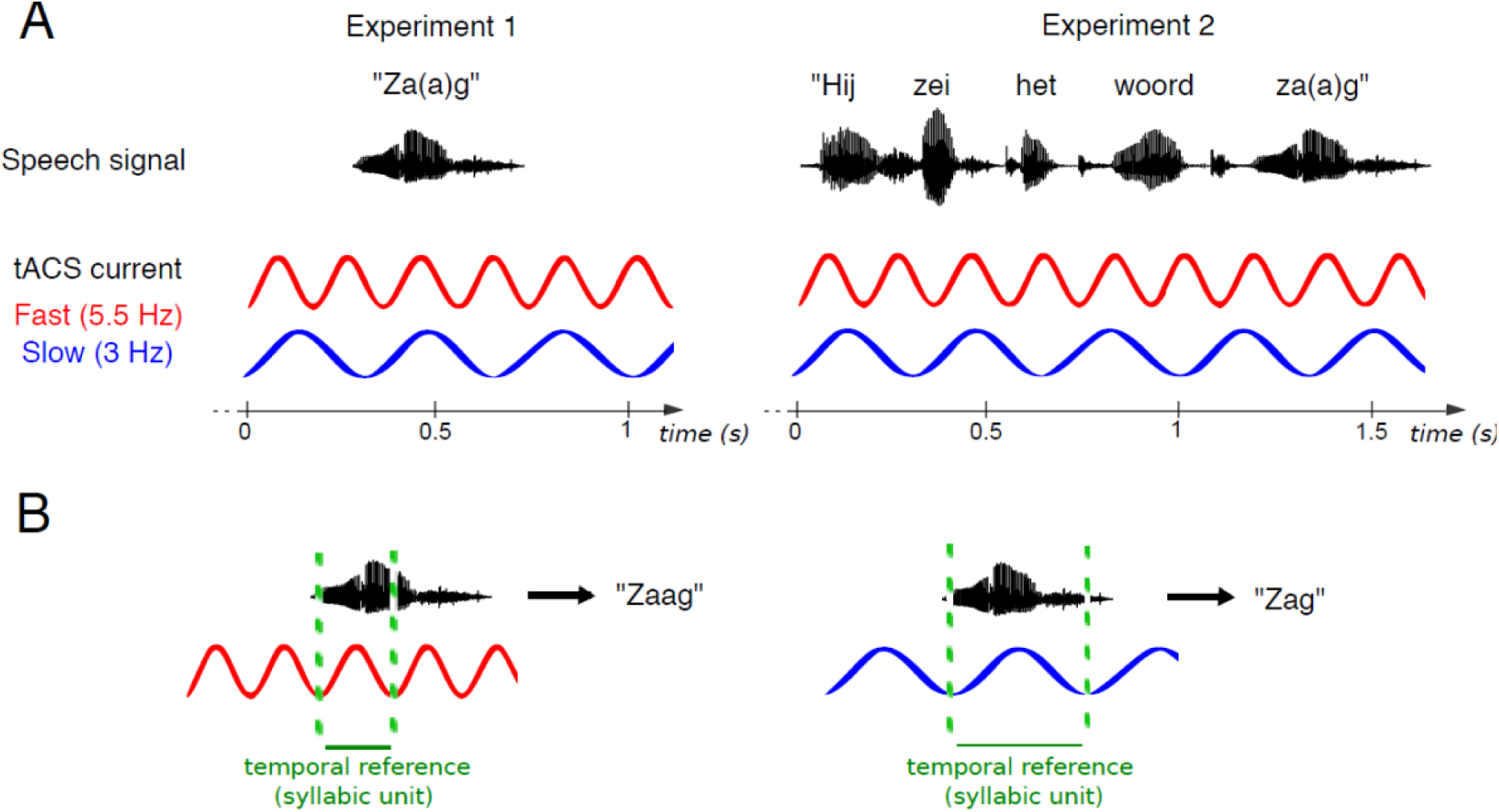
Experimental design and predictions. A) Participants listened to Dutch words that contained an ambiguous vowel (short “a” (/α/) – long “aa” (/a:/) contrast). The two vowels are dissociable based on both duration and spectral properties (2nd formant frequency, F2). Based on the perceived vowel, the words could be perceived as two distinct Dutch words with different meanings (e.g., “zag”, *saw [verb]* vs. “zaag”, *saw [noun]*). While participants listened to these words in isolation (Experiment 1) or in a sentence with a 4 Hz syllabic rate (Experiment 2), we applied continuous tACS above their auditory cortices at different frequencies (3 Hz and 5.5 Hz). B) We predicted that tACS entrains oscillations that act as temporal references for speech parsing. The change in frequency would bias the perceived duration of the chunked syllabic units and their constitutive phonological segments. More specifically it would bias the perceived duration of the ambiguous vowel (duration overestimation under Fast tACS, evidenced by a greater proportion of long vowel percepts; and underestimation under Slow tACS, with a lower proportion of long vowel percepts), leading to the perception of different words.

In two experiments, we asked Dutch participants to listen to Dutch words that contained a vowel which was ambiguous with regards to its duration (short “a”, /ɑ/ – long “aa”, /aː/ contrast). The words could be perceived as two distinct Dutch words with radically different meanings (e.g. “tak”, *branch*; “taak”, *task*). While participants listened to speech, we applied continuous tACS stimulation above the auditory cortices at different frequencies (3 Hz and 5.5 Hz) (Fig. 1A). We expected that tACS stimulation at different entrainment frequencies would entrain corresponding neural oscillations, and that these oscillations would influence temporal predictions, as reflected in how the words are perceived. Specifically, we predicted that stimulating the brain at a tACS frequency faster than the speech syllabic rate would lead to an overestimation of the speech segments’ duration (and in particular of the ambiguous vowel), inducing a greater proportion of long vowel percepts; conversely stimulating at a slower tACS frequency would lead to underestimation of the vowel duration (Fig. 1B), and fewer long vowel percepts.

## Experiment 1

### Methods

#### Participants

Twenty-five native Dutch participants (mean age: 23, 17 females) took part in the study. All participants were suited to undergo non-invasive brain stimulation as assessed by prior screening. They reported no history of neurological or hearing disorders, and gave their written informed consent before taking part in the study. One participant was excluded during tACS preparation due to intolerance to the electric stimulation. Another participant’s data were excluded due to a recording error. In total, data from 23 participants remained for analysis. The experimental procedure was approved by the local ethics committee (Ethical Review Committee Psychology and Neuroscience, Maastricht University).

#### Auditory stimuli

The speech stimuli were a subset of words previously used in (Kösem et al. 2018). A female native speaker of Dutch produced nine Dutch word pairs that only differed in their vowel, for instance, ‘‘zag’’ *(saw [verb])* vs. “zaag” (*saw [noun])*. The vowels for each word were constructed by selecting one long vowel “a” (/a:/) and manipulating its spectral and temporal properties, since the Dutch “a” (/ɑ/) – “aa” (/a:/) contrast is cued by both spectral and temporal characteristics (Audio S1 and S2) (Bosker 2017; Kösem et al. 2018). The temporal manipulation involved compressing the vowel to a duration of 140 ms using PSOLA in Praat (i.e., maintaining the original pitch contour) (Boersma and Weenink 2007). Spectral manipulations were based on Burg’s LPC method in Praat, with the source and filter models estimated automatically from the selected vowel. The formant values in the filter models were adjusted to result in a constant F1 value (740 Hz, ambiguous between “a” and “aa”) and 13 different F2 values (1100-1700 Hz in steps of 50 Hz). Then, the source and filter models were recombined and the new vowels were adjusted to have the same overall amplitude as the original vowel. This manipulation procedure resulted in a vowel with an ambiguous duration, but with spectral properties spanning a continuum from “a” and “aa”. Finally, the manipulated vowel tokens were combined with one consonantal frame (e.g., /z_x/) for each of the nine word pairs.

#### tACS settings

The tACS montage followed the montage used by (Riecke et al. 2015) to stimulate the auditory cortices. Square rubber electrodes were attached to the scalp with conductive adhesive paste at positions defined by the International 10-20 system. Two electrodes were placed over the temporal cortices (centered on positions T7 and T8) and two other electrodes were placed symmetrically to the left and right side of the midline (respectively) so that their long sides were centered on the vertex (position Cz) and bordering each other. A sinusoidal current with fixed starting phase was applied to the circuit above each cerebral hemisphere using two battery-operated stimulator systems (Neuroconn, Ilmenau, Germany). To create two approximately equivalent circuits, the skin was prepared so that the impedances of the left-lateralized and right-lateralized circuit were matched while keeping the net impedance below 10 kΩ (left: 3.8 ± 1.8 kΩ, right: 3.7 ± 1.8 kΩ, mean ± SD). The sinusoidal current was presented at two frequencies: 3 Hz and 5.5 Hz. The choice of these frequencies was based on the related previous MEG speech study (Kösem et al. 2018). Prior to the main experiment, tACS intensity was set individually by reducing the peak amplitude of the current simultaneously for both circuits in 0.1-mA steps from 1 mA to the point where participants reported feeling comfortable or uncertain about the presence of tACS under every electrode (on average 0.9 ± 0.1 mA, mean ± SD across participants).

For each tACS run of the experiment, tACS was continuously applied and its amplitude was ramped up over the first 10 s of the run using raised-cosine ramps during which no trials were presented. For runs comprising sham stimulation, this onset ramp was followed by an additional offset ramp lasting 30 s. Ramps at the end of the run were flipped, that is, they followed the reverse trajectory. Prior to the experiment, three waveforms were generated individually for each run (sampling rate: 16.5 kHz) that defined the acoustic stimulation, the electric stimulation, and the onsets of experimental trials (trial triggers) within the entire run, respectively. During the experiment, each of these waveforms was continuously fed in chunks into a separate channel of a digital-to-analog converter (National Instruments) operated by Datastreamer software (ten Oever et al. 2016). The outputs of the two ‘stimulation channels’ were further split and fed into stimulation devices (stereo soundcard and two tACS systems; see previous two sections). The ‘trigger channel’ output was fed into a PC on which Presentation software was running to control visual stimulation and button response acquisition.

#### Procedure

Participants were first familiarized with the auditory stimuli and task. They were presented with a vowel categorization task to estimate individual perceptual boundaries between “a” and “aa”. This pretest involved the presentation of the target word ‘‘dat’’, *that* - ‘‘daad’’, *deed* in isolation with 13 different equidistant F2 values between 1100 and 1700 Hz (with 9 repetitions of each F2 value). The F2 values were presented in random order. Participants were asked to listen to the spoken words while fixating a fixation cross on the screen with the two response options presented left and right (‘‘a’’ or ‘‘aa’’; position counter-balanced across participants), and to report what vowel they heard by pressing a button after each word presentation. Based on this pretest, individual psychometric functions were determined and the three F2 values yielding the 25%, 50%, and 75% long vowel “aa” categorization points were selected for the main experiment. This meant that the vowels spanned an ambiguous range, potentially allowing for the largest biasing effects, while at the same time providing participants sufficient variation to make the categorization task feasible.

The main experiment consisted of five 10 min-long runs (two runs with 3 Hz tACS; two runs with 5.5 Hz tACS; and one sham run) with short breaks in between. Each run contained 162 trials. Participants were asked to perform the same vowel categorization task as in the pretest, but this time all word pairs were presented. Subjects were blinded for stimulation conditions and runs were presented in random order. The sham run was identical to the tACS runs, except that it involved no electric stimulation beyond the on/off ramps (see tACS Settings section). In the stimulation conditions, the onsets of the target words appeared at six different phases of the tACS current (30, 90, 150, 210, 270 and 330°). During debriefing, participants were asked to provide a percentage for each run quantifying their confidence that they received electric stimulation. Participants’ confidence reports did not significantly differ between stimulation runs versus sham runs (*t*22 = −0.8, *p* = 0.42), suggesting that they were unaware of whether they received stimulation or sham stimulation.

#### Data analysis

We analyzed the effect of tACS Frequency condition (Fast: 5.5 Hz; Slow: 3 Hz) and Sham stimulation on the proportions of long vowel “aa” responses. Trials containing no button response (3.1 ± 8.5 % of all trials, mean ± SD across participants) and trials presented during Sham on-/off-ramps were discarded from the data analysis. Statistics were performed using Generalized Linear Mixed Models (GLMM) (Quené and van den Bergh 2008) with a logistic linking function as implemented in the lme4 library (version 1.0.5) (Bates et al. 2015) in R (Team 2013). Data were analyzed for fixed effects of Vowel F2, TACS Frequency, and their interaction. The model mapped the fast tACS Frequency onto the intercept, testing two contrasts: fast vs. slow, and fast vs. sham. The last contrast slow vs. sham was tested using a mathematically equivalent model with the same log-likelihood, achieved by mapping the slow tACS Frequency onto the intercept. The model included random intercepts for Participants and Word Pair, with by-participant and by-word pair random slopes for Vowel F2 (Barr et al. 2013). More complex random-effects structures failed to converge.

Supplementary phase analyses were performed by reconstructing a time series (composed of the six tACS phases at which the target word was presented) for each stimulation condition. The phase that most effectively biases perception may vary across individuals due to individual differences in anatomy. To compensate for such possible inter-individual variations, the maximum of the reconstructed series was aligned to the phase associated with strongest long /a:/ vowel percepts (labeled as phase 90°).

### Results

We expected that the rhythmic electric brain stimulation would entrain auditory cortices in a frequency-specific manner and hypothesized, based on (Kösem et al. 2018), that this would influence the perceived duration of the words’ vowels. Specifically, we expected to find a higher proportion of long “aa” responses in the 5.5 Hz tACS condition as compared to the 3 Hz tACS condition. However, against our expectations, no effect of TACS Frequency was observed (fast vs. slow: *p* = 0.460; fast vs. sham: *p* = .328; slow vs. sham: *p* = .686, Fig. 2). The proportion of long vowel responses was not significantly different for the Fast tACS frequency condition than for the Slow tACS frequency condition

**Figure 2:**
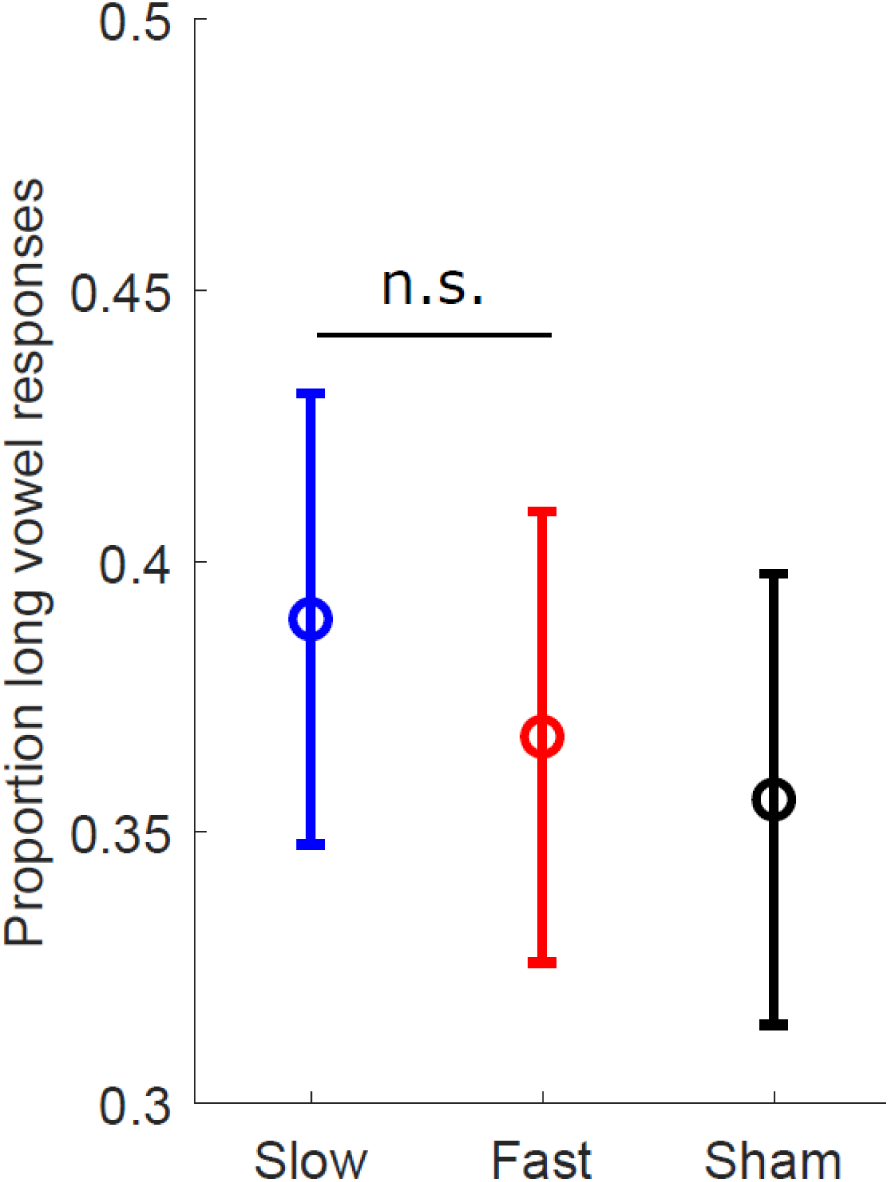
TACS frequency effects in experiment 1. TACS frequency did not significantly influence word perception. Proportion of long vowel word response during Slow (3 Hz) tACS (blue), Fast (5.5 Hz) tACS (red), and Sham stimulation (black) pooled across vowel F2s. No significant effect of stimulation frequency was found. Bars denote s.e.m.

To control that participants paid attention to the task and relied on acoustic cues to provide their response, we presented vowels with three distinct F2 frequencies (one ambiguous F2 value, one F2 value biasing participant reports toward short /α/ responses, and one F2 value biasing participant reports toward long /a:/ responses). The results suggest that participants indeed paid attention to the stimuli as they relied on the spectral cues to categorize the vowels: the vowel F2 had an effect on target word perception (*β* = 0.879, *SE* = 0.177, *z =* 4.957, *p* < .001), indicating that vowels with higher F2 were more likely to be perceived as long “aa”,. Partial evidence was found for interactions between Vowel F2 and tACS Frequency, but only for contrasts with the sham condition (Vowel F2 * the contrast between fast vs. sham: *β* = − 0.120, *SE* = 0.068, *z =* −1.764, *p* = .078; Vowel F2 * the contrast between slow vs. sham: *β* = − 0.136, *SE* = 0.673, *z =* −2.024, *p* = .043), meaning that perceptual differences between the two tACS conditions were not significantly different across F2 vowels.

We observed no significant effect of tACS phase on target word perception. Specifically, we analyzed perceptual reports for each tACS phase, after realignment to the phase associated with strongest long “aa” vowel percepts (see methods). Under the hypothesis that oscillatory phase modulates target word perception, we expected a bias toward long “aa” vowel percepts at phases neighboring the best phase whereas a bias towards short vowel word percepts should be observable at opposing phases. To test this prediction, long “aa” categorization proportions were averaged across the hypothesized positive half-wave (phases 30° and 150°; excluding 90°, which trivially represented the maximum value due to the phase realignment) and across the hypothesized negative half-wave (phases 210°, 270°, and 330°), and then the two resulting averages were statistically compared. Similar GLMMs as reported above were used, with the predictor Realigned Oscillation Half Cycle (positive half cycle coded as +0.5, negative half cycle as −0.5), which yielded no significant effect of oscillation half cycle (*p* = .15).

In sum, the results from Experiment 1 showed no significant influence of tACS frequency (or phase) on the perception of ambiguous words presented in isolation. A potential explanation for this null result is that low-frequency tACS effects on speech perception may be more readily observable when target words are presented in a (quasi-) rhythmic auditory context as in previous studies (Riecke et al. 2018, 2015; Wilsch et al. 2018), potentially because tACS may more strongly affect neural rhythms that are already present (Reato et al. 2010). In a second experiment, we tested if tACS influences speech perception when prior auditory input has already brought auditory cortices in an entrainment regime, by presenting ambiguous target words at the end of spoken sentences.

## Experiment 2

### Methods

#### Participants

31 native Dutch participants (mean age: 23, 18 female) took part in the study. All participants performed prior screening as in experiment 1. Two participants were excluded due to a bias in speech perception observed during the pretest (proportion of long “aa” words > 90%). One participant was excluded due to a recording error. In total, 28 participants remained for analysis.

#### Auditory stimuli

As in Experiment 1, the same female native speaker of Dutch produced nine Dutch word pairs that only differed in their vowel. In Experiment 2, these words were produced at the end of the fixed sentence frame “Hij zegt het woord [target]” *He says the word [target]* (Audio S3).

Target words were excised and manipulated to be ambiguous in vowel duration and quality. First, the durations of the two vowels of each pair were set to the mean vowel duration of that pair (*M* = 136 ms). Then, using sample-by-sample linear interpolation, we mixed the weighted sounds of the pair (11-point continuum; step 1 = 100% “a” + 0% “aa”; step 6 = 50% “a” + 50% “aa”; step 11 = 0% “a” + 100% “aa”; i.e., a step size of 10%) to create eleven different steps changing in vowel quality. We used this interpolation method because it resulted in more natural sounding output, while it also resulted in spectral vowel continua – similar to Experiment 1. These manipulated vowels were then spliced back into the consonantal frame from the “aa”-member of each pair, and concatenated onto one fixed token of the context sentence. This token of “Hij zegt het woord...” had a duration of 1100 ms and a pronounced peak at 4 Hz in its modulation spectrum, given the four monosyllabic words, falling in between the two tACS stimulation frequencies.

#### tACS settings

All tACS parameters were set as described for experiment 1. The average impedances of the left-lateralized and right-lateralized circuit were 5.3 ± 2.2 kΩ and 5.4 ± 2.4 kΩ, respectively, and the average tACS intensity was 0.9 ± 0.1 mA as before (mean ± SD across participants).

#### Procedure

The second experiment consisted of two acquisition sessions because of the increased duration of trials in comparison to experiment 1 (as full sentences were presented). In the first acquisition session, participants were familiarized with the stimuli with a vowel categorization task as in experiment 1. Each session consisted of six 7.5-minute-long runs (four stimulation runs and two sham runs) of 81 trials with short breaks in between. As in experiment 1, participants were asked to listen to the sentences and report their perception of the last word.

The onsets of the target words appeared at six different phases of the tACS current. Participants’ confidence reports did not significantly differ between tACS runs versus sham runs (*t*_27_ = 0.1, p = 0.92).

#### Data analysis

The same analyses were performed as in Experiment 1. Trials containing no button response (1.4 ± 4.0% of all trials, mean ± SD across participants) and sham trials presented during tACS on-off ramps were discarded from the data analysis. A GLMM and was used to test for fixed effects of Vowel F2 and tACS Frequency. Adding the interaction term did not improve model fit, as evidenced by log-likelihood model comparison (*p* = .184). The model also included random intercepts for Participants and Target Pair, with by-participant and by-word pair random slopes for Vowel F2.

#### Results

As in Experiment 1, the vowel F2 had an effect on target word perception F2 (*β* = 2.960, *SE* = 0.359, *z =* 8.236, *p* < .001). In contrast with experiment 1, and in line with our hypothesis, the difference between Fast and Slow tACS frequency conditions was significant: 5.5 Hz tACS led to a small increase in the proportion of long vowel responses relative to 3 Hz tACS reliably across participants (fast vs. slow: *β* = −0.085, *SE* = 0.043, *z =* −1.979, *p* = .048, Fig. 3). Contrasts with the sham condition yielded no significant result (fast vs. sham: *p* = 0.475; slow vs. sham: *p* = .230), and no effect of tACS phase was observed (that is, after phase realignment, GLMMs with the predictor Realigned Oscillation Half Cycle showed no significant effect of half cycle; *p* = .52).

**Figure 3:**
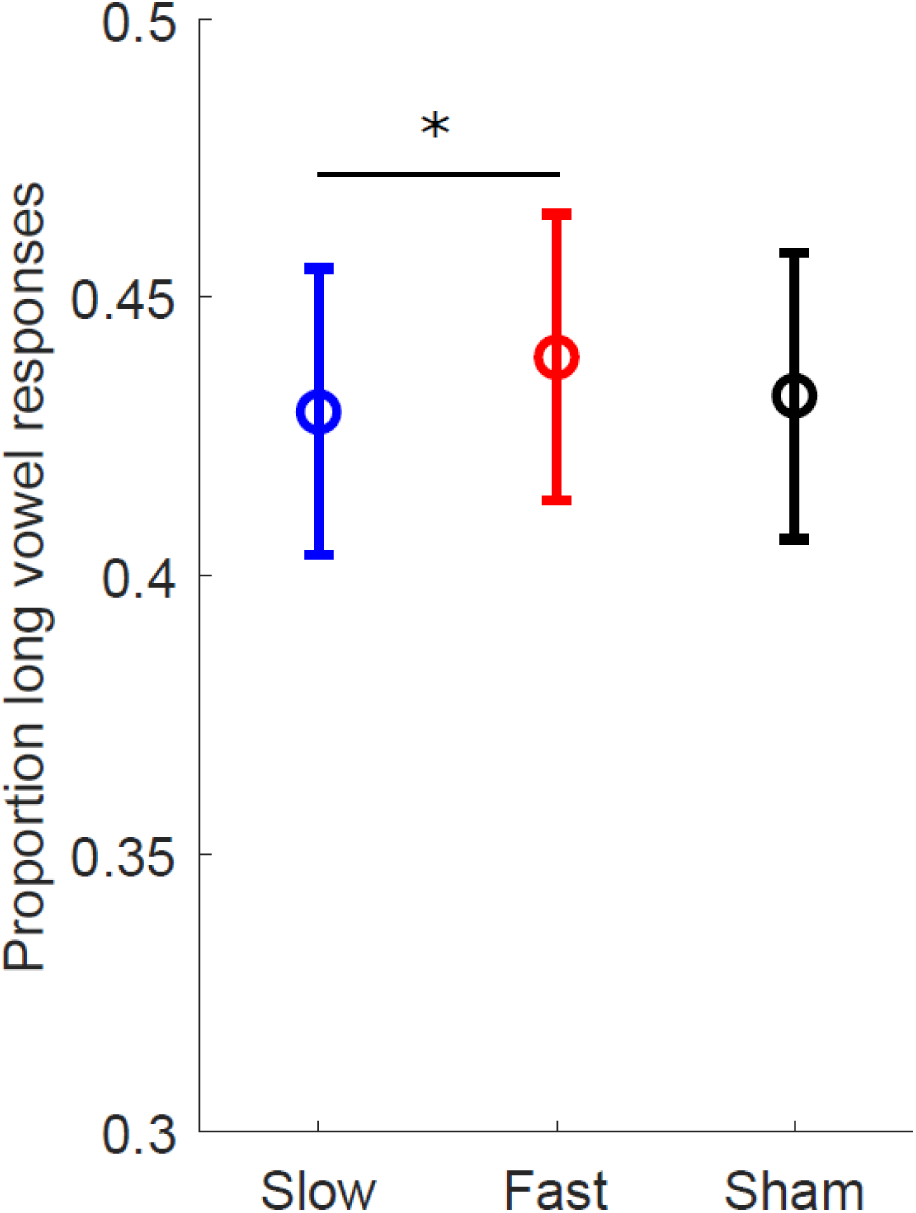
TACS frequency influenced word perception in Experiment 2. A) Percentage of long vowel word response during Slow (3 Hz) tACS (blue), Fast (5.5 Hz) tACS (red), and Sham stimulation (black). Bars denote s.e.m. * represents P<0.05.

Because we observed an effect of tACS Frequency between the fast and slow stimulation frequency conditions in Experiment 2 but not in Experiment 1, we additionally ran an omnibus analysis on the complete data set from both experiments. This omnibus GLMM was identical to the GLMM reported above, except that it additionally contained the fixed effect Experiment and an interaction term for tACS Frequency * Experiment. Adding this interaction term significantly improved model fit, as evidenced by log-likelihood model comparison (*χ2*(2) = 18.953, *p* < 0.001), and the two-way interaction was indeed significant for the fast vs. slow contrast (*β* = −0.209, *SE* = 0.060, *z =* −3.474, *p* < .001, Fig. 4). This indicates that the observed difference in perception between fast and slow tACS conditions was significantly larger in Experiment 2 compared to Experiment 1, where no statistically significant effect was found.

**Fig. 4:**
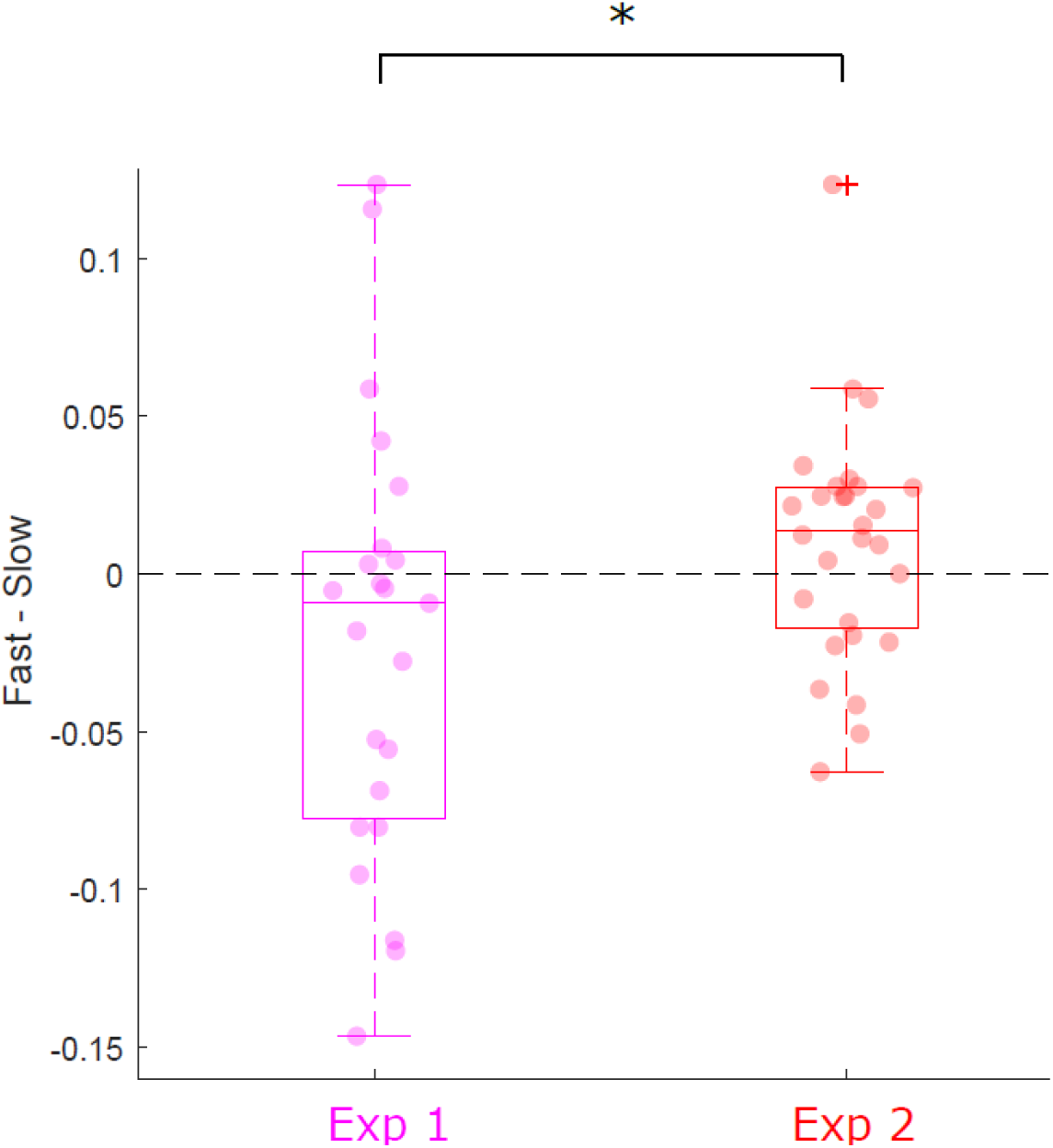
tACS frequency influences on speech perception is different across experiments. Box plots represent the distribution of the difference between Fast and Slow tACS conditions in experiments 1 (*n* = 23) and 2 (*n* = 28). Each dot represents one participant. tACS Frequency has a larger effect on the perception of spoken words when they are presented in continuous speech (Experiment 2) vs. in isolation (Experiment 1). The central mark of the boxplot represents the median of the distribution, the edges of the box are the 25th and 75th percentiles, the whiskers extend to the most non-outliers extreme data points, and the cross represents and outlier. * represents P<0.05.

Considering that single words were presented in Experiment 1, while full sentences were presented in Experiment 2, these results suggest that tACS frequency effects on speech perception are more readily observable when target words are presented in a (quasi-)rhythmic auditory context.

## General Discussion

We tested the effect of tACS frequency (within the theta range) on the perception of speech content, following recent evidence suggesting that low-frequency neural entrainment to the speech envelope influences the categorization of phonemes and therefore the perception of words (Ten Oever and Sack 2015; Kösem et al. 2018). Our first experiment showed no significant effect of tACS frequency on word perception. Based on previous tACS studies on the perception of continuous speech (Wilsch et al. 2018; Riecke et al. 2018), we reasoned that this null result may reflect the use of isolated words. Therefore, we further hypothesized that tACS frequency effects on perceptual speech segmentation require the speech to be presented in a continuous (quasi-)rhythmic auditory context. Our second experiment provides support for our hypotheses: we observed that tACS presented at a fast frequency elicits on average more long vowel percepts than tACS presented at a slower frequency, consistent with the idea that entrainment of faster neural oscillations results in a denser sampling of speech input (Kösem et al. 2018). We additionally found a significant difference with respect to the tACS frequency effect on speech segmentation across experiments: in line with our secondary hypothesis, tACS frequency had a significantly larger influence on the segmentation of speech when the latter was presented in a continuous sentential context rather than as an isolated word.

These results suggest that tACS causally influences the perception of speech sounds. We interpret the outcomes as an indication that tACS influenced neural entrainment, which reflect a neural mechanism by which the input speech signal is sampled at the appropriate temporal granularity (Giraud and Poeppel 2012; Ghitza 2011). We used a tACS montage targeting auditory cortices (Riecke, Sack, and Schroeder 2015; Riecke et al. 2015), suggesting that the observed effect occurs in auditory cortical areas involved in speech processing. This notion is corroborated by findings showing that phonological information may be decoded from early auditory oscillatory activity (Di Liberto, O’Sullivan, and Lalor 2015; Ten Oever and Sack 2015), and that behavioral perceptual biases induced by fast vs. slow speech rhythms arise early in perception (Maslowski, Meyer, and Bosker 2019) and independently from attention (Bosker, Reinisch, and Sjerps 2017). Our results show no significant effect of tACS phase on vowel perception. Although not the focus of our study, this absence of a phase effect in the presence of a frequency effect is in line with previous results from a speech study that used auditory, instead of electric, stimulation to manipulate neural entrainment (Bosker & Kösem, 2017). It contradicts phase effects observed in a previous tACS study that investigated intelligibility of continuous speech in noise (Riecke et al., 2018), suggesting that such phase effects arise during behavioral tasks that require processes related to auditory stream segregation.

The combined outcomes suggest that tACS may modulate the perceptual sampling of speech more effectively in the context of continuous speech than for single word presentations. A tentative interpretation for our results is that tACS may only have a modulatory influence on brain oscillations that have already been entrained by prior sensory input. That is, tACS at the relatively weak stimulation intensity used here (~1.8 mA peak-to-peak) may be more effective in modulating a pre-existing neural entrainment (induced by a given rhythmic sensory input) than in inducing neural entrainment in the absence of external sensory rhythms. Concurrent recordings of neural activity during transcranial stimulation show that weak-intensity tACS may not induce neural oscillations when neural activity is not strongly rhythmic (Lafon et al. 2017), but could affect already present narrow-band neural rhythms (Reato et al. 2010). This could explain why low-frequency tACS is most effective at frequencies close to ongoing brain rhythms (Kanai et al. 2008), and in sensory stimulus-induced entrainment settings (Riecke et al. 2015, 2018; Zoefel, Archer-Boyd, and Davis 2018; Wilsch et al. 2018). We speculate that in Experiment 2, tACS at 3 and 5.5 Hz modulated the frequency of neural oscillations that were entrained to the envelope of the continuous speech stimuli, which fluctuated most strongly at 4 Hz. When words were presented in isolation there was no rhythmic auditory stimulation to entrain neural oscillations, and as such tACS probably had less influence on the brain processes that involve entrained oscillations, such as temporal predictions (Stefanics et al. 2010; Kösem et al. 2018).

Alternatively, tACS may have affected word perception differently across our two experiments because neural responses to the target word differed when it was presented in continuous speech as compared to when it was presented in isolation. Neural responses to a word are likely attenuated in continuous speech, considering that the response evoked by an acoustic input reduces when the input is preceded by a temporally regular sequence of stimuli (Todorovic and de Lange 2012; Costa-Faidella et al. 2011). Moreover, tACS-induced periodic alterations in neural excitability may affect sensory stimulus processing most effectively when the stimuli are near threshold. Therefore, tACS probably modulated neural activity in our two experiments in a similar fashion, but this modulation was stronger in Experiment 2 as neural responses evoked by the target word were weaker and thus more susceptible to tACS-induced modulations.

The results therefore support oscillatory models of speech processing in certain contexts, i.e. during continuous speech listening. However, the size of the effect in Experiment 2 – although statistically significant – was rather modest. Our experiments involved a relatively large number of participants, multiple sessions, many repetitions of the same stimuli, and ambiguous speech sounds that are most sensitive to perceptual biases. As such, the present outcomes do not warrant bold claims about the alleged ‘brain hacking’ potential of transcranial electrical brain stimulation. In fact, concerns have been expressed recently about the efficacy of tDCS and tACS in directly modulating neural activity and behavior, in particular when applied currents are weak (~1-2 mA) (Liu et al. 2018; Opitz et al. 2016). At this current strength, effects on neural activity are observable, but may be restricted to temporal biasing of spikes and/or modulation of ongoing neural rhythms of similar frequency as the applied current (Liu et al. 2018; Krause et al. 2019). Our behavioral findings fit with these observations and point to an interesting role of sensory stimulation history on tACS efficacy, which should inspire further investigation into the constraints under which tACS modulates human behavior, and speech comprehension in particular.

## Acknowledgements

This study was supported by the Netherlands Organization for Scientific Research (NWO) Gravitation Grant 024.001.006 awarded to the Language in Interaction Consortium, a Marie Sklodowska-Curie Individual Fellowship (grant number 843088) to AK, a James S. McDonnell Foundation Understanding Human Cognition Collaborative Award (grant number 220020448) and Welcome Trust Investigator Award in Science (grant number 207550) to OJ. We would like to thank Annelies van Wijngaarden for the recordings of her voice.

